# Multi-omic analyses reveal antibody-dependent natural killer cell-mediated cytotoxicity in autoimmune thyroid diseases

**DOI:** 10.1101/662957

**Authors:** Tiphaine C. Martin, Kristina M. Illieva, Alessia Visconti, Michelle Beaumont, Steven J. Kiddle, Richard J.B. Dobson, Massimo Mangino, Ee Mun Lim, Marija Pezer, Claire J. Steves, Jordana T. Bell, Scott G. Wilson, Gordan Lauc, Mario Roederer, John P. Walsh, Tim D. Spector, Sophia N. Karagiannis

**Affiliations:** Department of Twin Research and Genetic Epidemiology, King’s College, London, United Kingdom; School of Biomedical Sciences, University of Western Australia, Crawley, Western Australia, Australia; Department of Oncological Sciences, Icahn School of Medicine at Mount Sinai, New York City, NY 10029, USA; Tisch Institute, Icahn School of Medicine at Mount Sinai, New York City, NY 10029, USA; St John’s Institute of Dermatology, School of Basic & Medical Biosciences, King’s College, London, Guy’s Hospital, London, United Kingdom; Breast Cancer Now Research Unit, School of Cancer & Pharmaceutical Sciences, King’s College London, Guy’s Cancer Centre, London, United Kingdom; Department of Biostatistics and Health Informatics, Institute of Psychiatry, Psychology and Neuroscience, King’s College, London, United Kingdom; MRC Biostatistics Unit, University of Cambridge, Cambridge, CB2 0SR, UK; Health Data Research UK (HDR UK), London Institute of Health Informatics, University College London, London, United Kingdom; NIHR Biomedical Research Centre at Guy’s and St. Thomas’s NHS Foundation Trust, London, United Kingdom; Department of Endocrinology and Diabetes, Sir Charles Gairdner Hospital, Nedlands, Western Australia, Australia; Medical School, The University of Western Australia, Crawley, Western Australia, Australia; PathWest Laboratory Medicine, QEII Medical Centre, Nedlands, Western Australia; Faculty of Pharmacy and Biochemistry, University of Zagreb, Zagreb, Croatia; Genos, Glycoscience Research Laboratory, Zagreb, Croatia; ImmunoTechnology Section, Vaccine Research Center, NIAID, NIH, Bethesda, MD 20892, USA

**Author notes:** Equal contribution as a joint senior (Tim D. Spector, Sophia N. Karagiannis). **Author Contributions:** T.C.M. designed the general study, developed, applied the biostatistical analysis, interpreted findings, drafted the manuscript and produced the figures and tables. T.D.S. coordinated the recruitment of the TwinsUK cohort and research projects applied to this cohort. M.P. and G.L. performed glycan assays in the TwinsUK cohorts. S.J.K, R.J.B.D and C.S. coordinated proteomic assays in the TwinsUK cohort. M.M coordinated the detection of immune cell traits in the TwinsUK cohorts and performed GWASs on these immune cell traits. M.R. performed the detection of immune cell traits in the TwinsUK cohorts. A.V. removed the batch effect on immune cell traits. J.P.W. and S.G.W coordinated, analyzed and interpreted thyroid function tests in the TwinsUK cohort. E.M.L. performed thyroid function tests in the TwinsUK cohort. M.B and J.T.B provided feedback on the paper. T.C.M., K.I., and S.N.K. edited the manuscript with input from all authors. All authors read and approved the final manuscript.

**Keywords:** Glycosylation, immune cell traits, proteomics, IgG, autoimmune thyroid diseases, Hashimoto’s disease, Graves’ disease, TPOAb, genetic variants, apoptosis, ADCC

## Abstract

The pathogenesis of autoimmune thyroid diseases (AITD) is poorly understood. We previously observed systemic depletion of IgG core fucosylation and antennary α1,2 fucosylation of peripheral blood mononuclear cells in AITD, correlated with thyroid peroxidase antibody (TPOAb) levels. We hypothesized that deficiency in IgG core fucose enhances antibody-dependent cell-mediated cytotoxicity of thyrocytes by TPOAb, contributing to thyroid autoimmunity. Multi-omic evaluations in 622 individuals (172 with AITD) from the TwinsUK cohort showed decreased IgG core fucosylation levels associated with a subpopulation of natural killer (NK) cells featuring CD335, CD314, and CD158b immunoreceptors, and increased levels of apoptosis-associated Caspase-2 and Interleukin-1α, positively associated with AITD. AITD-associated genetic variants rs1521 and rs3094228 alter expression of thyrocyte ligands of the CD314 and CD158b immunoreceptors on NK cells. The combination of low-core fucose IgG associated with an NK cell subpopulation and genetic variant-promoted ligand activation in thyrocytes may promote antibody-dependent NK cell-mediated cytotoxicity of thyrocytes in AITD.

## Introduction

Autoimmune thyroid diseases (AITD) are a class of chronic, organ-specific autoimmune disorders that affect the thyroid gland with a high genetic heritability (55-75%)^1–4^. AITD are diagnosed in approximately 5% of the European population with a gender disparity (i.e., women: 5-15%; men: 1-5%), and represent the most common group of autoimmune diseases^5–7^. AITD are characterised by a dysregulation of the immune system affecting several biological structures and processes, such as antigen/antibody/effector cell complex formation^8–10^, likely driven by a combination of genetic and environmental factors^11^.

Pathologically, AITD are characterized by the production of autoantibodies against the three key thyroid proteins (thyroid peroxidase (TPO), thyroglobulin (Tg), and the thyroid-stimulating hormone (TSH) receptor (TSH-R)), infiltration of the thyroid gland by immune cells (e.g. lymphocytes, NK cells, monocytes, and macrophages), as well as the formation of germinal centers in the thyroid gland^12^. Additionally, several vital pathological players and biomarkers can be found in peripheral blood, such as high levels of thyroid autoantibodies and dysregulated TSH levels^13, 14^. However, some controversy exists surrounding the immune cell composition in peripheral blood associated with AITD. Some studies have failed to observe a significant difference in peripheral blood immune cell composition between AITD patients and healthy individuals^15^, whereas others have reported significant differences in particular cell types or specific marker expression on immune cells.^16^. Immune cells, thyroid autoantibodies, and secreted proteins including cytokines may play critical roles in AITD development^17^ and could participate in immune responses including antibody-dependent cell-mediated cytotoxicity (ADCC) pathways^18, 19^.

ADCC is triggered with the formation of antigen/antibody/Fc receptor complexes bringing the effector cell (macrophages, NK cells) and the target cell (expressing the antigen) in close contact. As the formation and the function of the antigen/antibody complex are modulated by post-translational modifications of proteins^20, 21^, we previously studied the glycosylation of total immunoglobulin G (IgG) and peripheral blood mononuclear cells (PBMC) in patients with AITD^4^. We identified a depletion of IgG core fucosylation and antennary α1,2 fucosylation of PBMCs in peripheral blood, associated with TPOAb levels and AITD status^4^. IgG core fucose, which is observed in approximately 95% of IgG in healthy individuals, is considered a “safe switch” by attenuating potentially harmful ADCC activity against self-antigens^22–26^. Therefore, it is possible that the deficiency of IgG core fucose observed in AITD and correlated with elevated TPOAb levels may be associated with TPOAb production in the thyroid gland and could be a player in autoimmune response in AITD by enhancing the cytotoxicity activities of TPOAb against thyrocytes through ADCC. Previous studies showed that afucosylated antibodies have a higher affinity (∼100-fold) in binding to the immunoglobulin receptor FcγRIIIa (CD16a), expressed on NK cells, macrophages and γδ T cells, and associated with enhanced ADCC potential *in vitro* and anti-tumor activity *in vivo*^22–24, 26^. Consequently, we hypothesized that particular immune features that accompany Fc-mediated functions of IgG, immune cells and associated inflammation could be detected in PBMC of patients with AITD.

Therefore, this *in silico* study aimed to: 1) investigate the association of different components of antigen/antibody/Fc receptor complexes with AITD, 2) identify interactions between these different immune components, and 3) elucidate genetic and environmental effects on these components within the bloodstream in 622 subjects from the TwinsUK cohort, of whom 172 have an AITD diagnosis. More precisely, we examined the association of total serum IgG glycosylation, immune traits, such as immune cell subpopulation frequencies (CSFs; i.e. the relative frequencies of circulating immune cell subsets), immune cell surface protein expression levels (SPELs; i.e. the measurement of the cell-surface expression of critical proteins) and secreted proteins, from the peripheral blood of patients with AITD compared with those in normal volunteer blood. Our study design is summarized in **Figure 1**, and the sample sizes of these different studies are described more in detail in **Supplement Table 1**.

**Figure 1.**
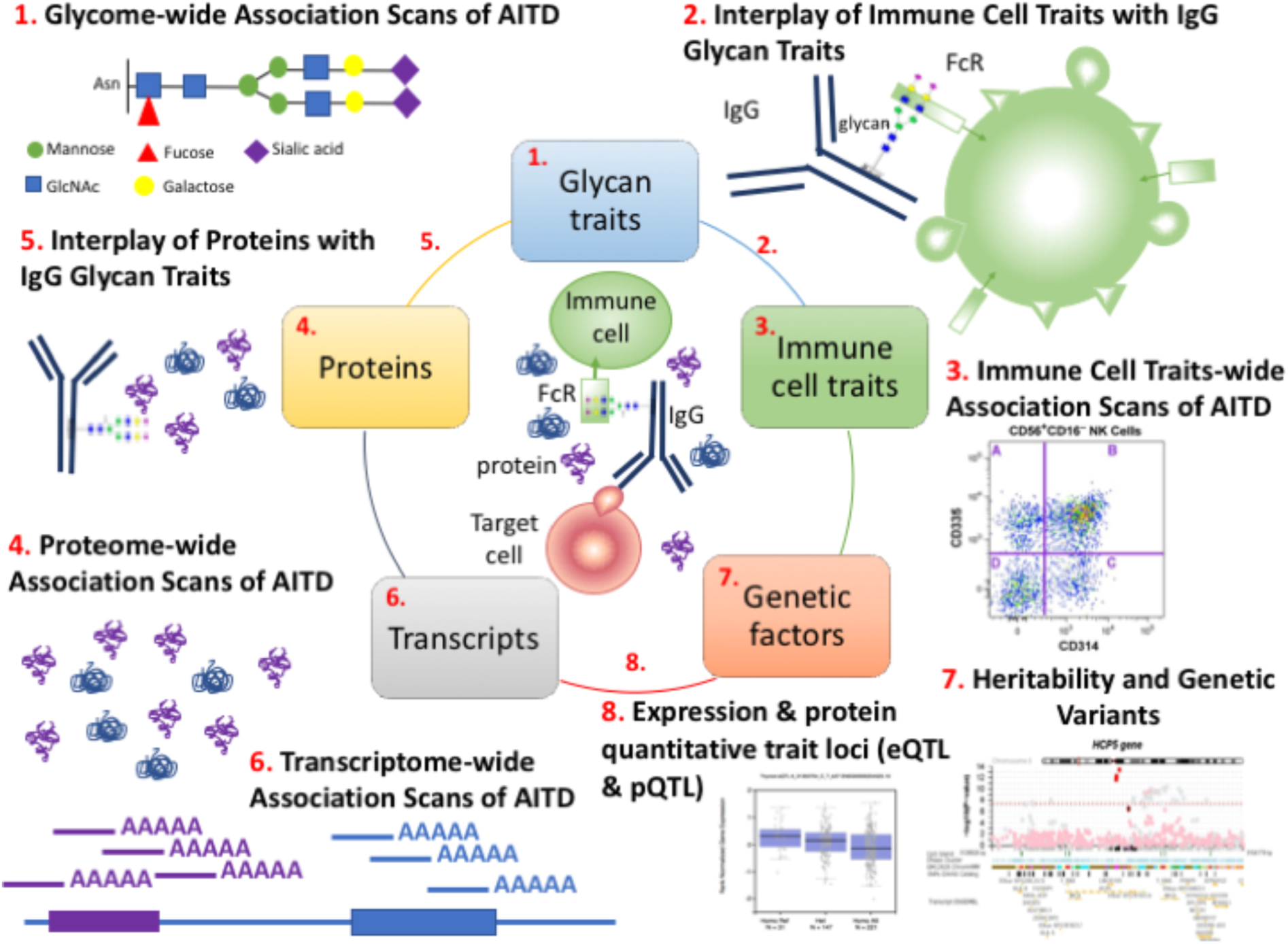
Multi-omics computational analyses used to study the components of antigen/antibody/effector cell complex structure in AITD. (1) We performed glycome-wide association studies of AITD and TPOAb levels in our previous work using 3,146 individuals from three European cohorts, including the TwinsUK cohort and identified 17 AITD-IgG N-glycan traits with seven replicated^4^. (2) Association study of total IgG N-glycan traits with 23,485 immune cell traits in 383 individuals from the TwinsUK cohort (regardless of disease status) showed that 6 out of the 17 AITD-IgG glycan traits were correlated with 51 immune cell traits featuring the CD335, CD134, and CD158b receptors. (3) However, none of these 51 immune cell traits appeared to be associated with AITD in up to 374 individuals. (4) We observed 3 out of 1,113 circulating proteins tested in plasma of almost 300 individuals shown to be associated with AITD status (TSH, Caspase-2, and Interleukin-1α). (5) Several secreted proteins were correlated with the level of plasma IgG glycan traits in 164 individuals, but none of them were also associated with AITD. (6) Although transcriptome-wide association studies of AITD, TPOAb level, and N-glycan structures were previously performed in approximately 300 individuals, no finding was significant in this study^10^. (7) Furthermore, we studied the genetic factors of these different findings. First, the heritability of secreted proteins was performed in this study whereas the heritability of AITD, TPOAb level and several omic features (IgG N-glycan traits and immune cell traits) were performed in previous studies of the TwinsUK cohort^4, 27–29^. No shared additive genetic variance between different phenotypes studied here (AITD status, TPOAb level, level of IgG N-glycan traits, of immune cell traits and of circulating proteins in the bloodstream) could be determined. (8) We identified genetic variants that alter the expression of genes, proteins and cell-bound immune receptors highlighted in this paper using the previous GWASs performed in the TwinsUK cohort or from GWAS catalog, eQTLs from GTEx project and pQTLs from INTERVAL project^27, 28, 30–33^. The sample sizes of these different studies are described in **Supplement Table 1**. GlcNAc = N-acetylglucosamine.

## Results

### Depletion of IgG core fucose is positively associated with increased CD158b^+^CD314^+^CD335^-^ NK cell subset count

IgG N-glycosylation is considered indispensable for the effector function of IgG and the control of inflammation^34–38^ and plays an essential role in the recognition and binding to Fc receptors of immune cells^37^. In our previous study^4^, we observed a deficiency of IgG core fucose in the peripheral blood of AITD patients, and it is known that IgG core fucose plays a role in ADCC^22–24, 26^. Recently, we performed genome-wide association studies (GWASs) on 78,000 immune cell traits using a high-resolution deep immunophenotyping flow cytometry analysis in 669 twins (497 discovery, 172 replication) from the TwinsUK cohort^27, 28^ and identified genetic variants known to confer autoimmune susceptibility^27, 28^. Following quality control of samples and immune cell traits, 383 individuals have been identified to have measurements of 23,485 immune cell traits and 17 AITD-IgG N-glycan traits. Consequently, we examined whether levels of the 17 IgG N-glycan traits previously associated with AITD (IGP2, IGP7, IGP8, IGP15, IGP21, IGP33, IGP36, IGP42, IGP45, IGP46, IGP48, IGP56, IGP58, IGP59, IGP60, IGP62 and IGP63) were also associated with the level of specific immune cell trait subpopulations in the TwinsUK cohort (**Supplement Table 1**). We carried out the association analyses taking into account immune trait correlations and the hierarchical nature between different immune cell traits as well as IgG N-glycan trait correlations. We identified 1,357 independent immune cell traits among 23,485 tested immune cell traits, 20 independent IgG N-glycan traits among 75 IgG N-glycan traits, and 6 independent AITD-IgG N-glycan traits among 17 AITD-IgG N-glycan traits using Li & Ji’s method^45^. Association studies of total IgG N-glycan traits with immune cell traits showed that 6 of the 17 significant IgG N-glycan traits (IGP2, IGP42, IGP46, IGP58, IGP59, IGP60) previously associated with TPOAb level and AITD status detected in the TwinsUK cohort, were also associated with 51 immune cell traits (**Supplement Table 2, Fig. 2)**. Three IgG N-glycan traits (IGP2, IGP42, IGP46) were negatively associated with the level of the activating subpopulation of CD16^+^CD56^+^CD158b^-^CD335^+^ NK cells and positively associated with the level of the CD16^+^CD56^+^CD335^-^ effector NK cell subpopulation and with the activating subpopulation CD16^+^CD56^+^CD158b^+^CD314+CD335^-^ NK cells^39–44^. In contrast, three other significant IgG N-glycan traits with core fucose (IGP58, IGP59, and IGP60) had the opposite effect associations with the same subpopulations of NK cells **(Fig. 2a)**. It is noteworthy that we previously showed that there are negative correlations between the set of IgG N-glycan traits without core fucose (IGP2, IGP42, IGP46) and the set of IgG N-glycan traits describing IgG core fucose (IGP58, IGP59, and IGP60)^4^ and these are highlighted in **Fig. 2b**. Moreover, we observed a strong correlation between these 51 immune cell traits (**Fig. 2c**). The presence of correlation patterns between the 17 AITD-IgG N-glycan traits (**Fig. 2b**) as well as between the 51 immune cell traits (**Fig. 2c**) is consistent with our observation of correlations between the 6 AITD-IgG N-glycan traits and the 51 immune cell traits **(Fig. 2a)**. Moreover, we extended our analysis to the 58 remaining IgG N-glycan traits also identified in our samples, but not associated with AITD, and we observed no significant association between them and the 23,485 immune cell traits.

**Figure 2.**
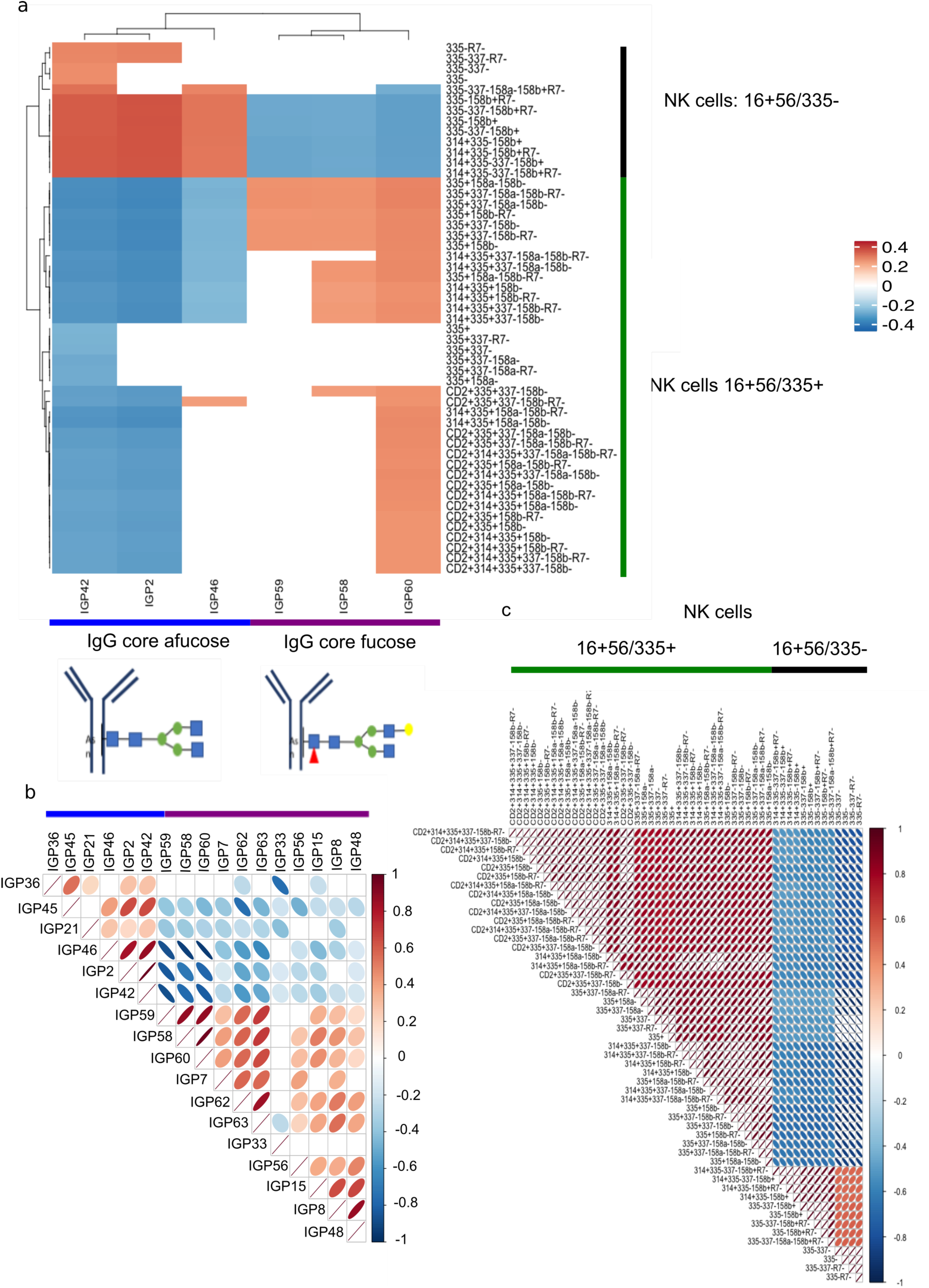
AITD-IgG N-glycan traits associated with a subpopulation of NK cells. a) Heatmap of immune cell traits associated with AITD-IgG N-glycan traits. The 51 NK cell types were significantly associated with 6 out of 17 AITD-IgG N-glycan traits previously identified^4^. Below the heatmap, there are one representative of IgG core afucose (IGP2) and one representative of IgG core fucose (IGP7), that were both associated with AITD and TPOAb levels^4^. b) Co-expressions between only 17 IgG N-glycan traits previously associated significantly with AITD status and TPOAb level^4^. c) Correlations between the profile of 51 immune cell traits that were associated significantly with at least one of 17 AITD-IgG N-glycan traits. The order of immune cell traits is the same as that in **Fig 2a**.

Consequently, we conclude that a subpopulation of NK cells with a combination of specific immunoreceptors (CD314, CD335, CD2, CD158a, CD158b, R7) is associated with the depletion of IgG core fucose observed in individuals with AITD.

### No significant association between immune cell traits and AITD or TPOAb level could be identified

We furthermore investigated whether the 23,485 peripheral blood immune cell traits are directly associated with TPOAb levels and AITD status in up to 374 individuals from the TwinsUK cohort (**Supplement Table 1**). We performed association analyses of TPOAb levels and AITD status with all 23,485 immune cell traits, with a particular focus on the 51 immune cell traits identified in the previous analysis (respectively 1,357 and 6 independent immune cell traits) (**Fig. 2c and 3a**). No significant associations of immune cell traits with TPOAb level or with AITD status could be detected (**Supplement Table 3**). These results suggest that even if there are correlations between IgG core fucose levels and immune cell traits in blood, and a correlation between IgG core fucose and AITD, the association between immune cell traits and AITD in blood appears to be mediated by more complex processes (**Supplement Fig. 1**).

**Figure 3.**
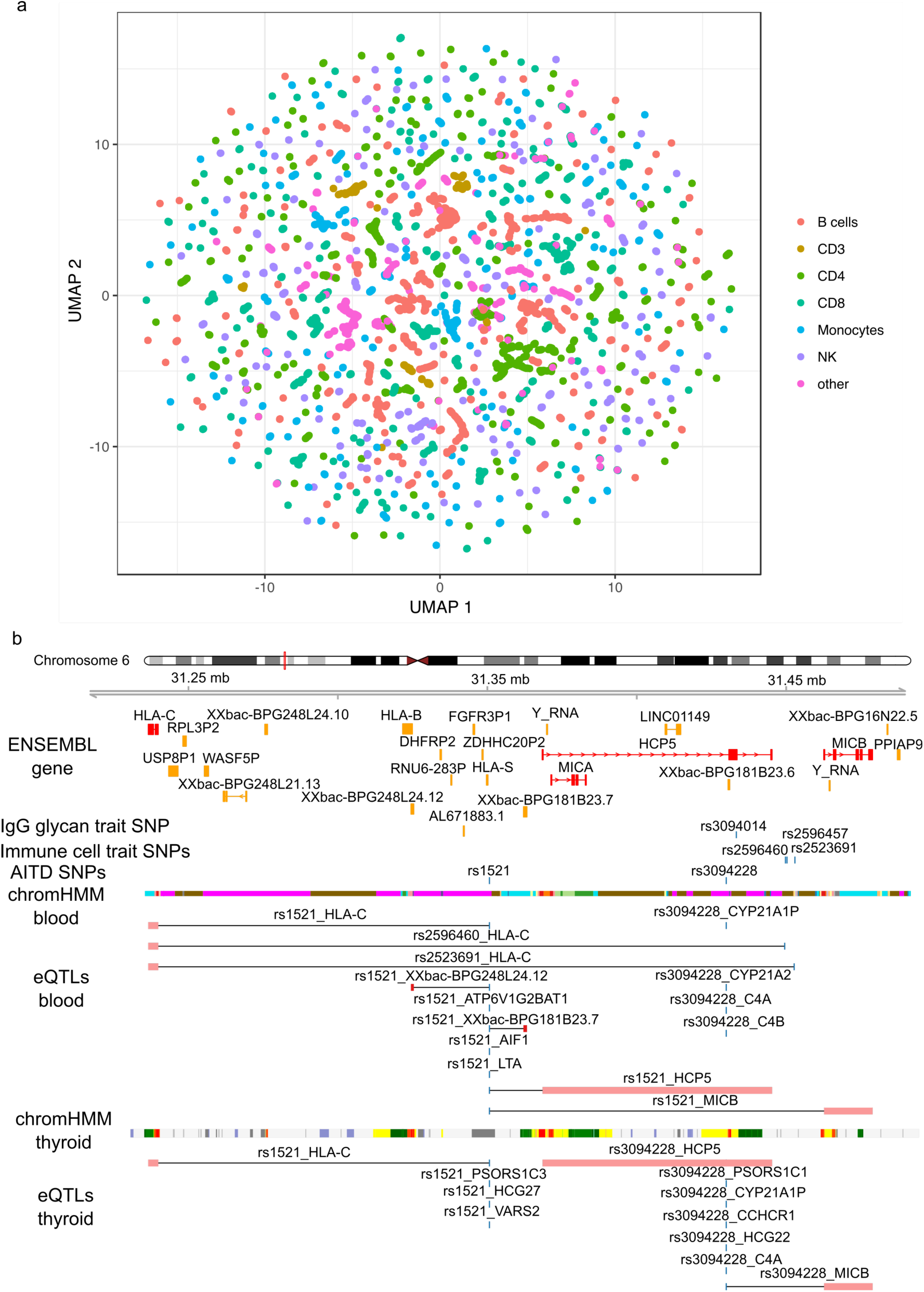
Association of immune cell traits with AITD status. a) Immune cell traits were arranged in two dimensions based on the similarity of their quantification profiles by the dimensionality reduction technique UMAP^50^ using R package umapr^51^. Some clusters that emerge spontaneously can be associated with specific immune cell types (colors). b) Annotation tracks around *MIC-A, MIC-B* and *HLA-C* genes visualize significant GWAS hits for immune cell traits, the ligands of certain immunoreceptors (such as NK), and thyroid phenotypes previously identified in the TwinsUK cohort as well as chromatin states identified using chromHMM from whole blood from ENCODE^52^ and thyroid cells from CEMT^53^ and eQTLs from GTEx project^32, 33^. The plot was produced using functions from R packages Gviz and coMET^54^.

### The genetic variants, rs1521 and rs3094228, previously associated with AITD alter the expression of the ligands of CD314 and CD158b immunoreceptors in the thyroid cells

Although overall little is known about the specific functions of NK cells based on the combination of their immunoreceptors, several studies describe the functions of each of three crucial NK receptors: CD335 (NKp46), CD314 (*NKG2D)* and the killer cell immunoglobulin-like receptors (KIRs) including CD158b. These are normally associated with activated NK cell states, T cell co-stimulation, and mediating tumor cell lysis^40, 42^. We thus inspected genetic variants associated with AITD, TPOAb levels, and immune cell traits from the previous GWAS^27, 28, 30^, and compared these with recent large-scale studies on tissue-specific expression quantitative traits (eQTLs), mainly from the GTEx project^32, 33^, to determine whether genetic factors could contribute to AITD, related immune features, or their pathways.

First, we identified loci and the reported genes with the lead single nucleotide polymorphisms (SNPs) associated with AITD as well as immune cell traits from previous GWASs^27, 28, 30^. No single lead genetic variants identified by our previous immune cell traits GWASs appeared to be also associated with at least one of thyroid phenotypes previously studied and published in the GWAS catalog (e.g., Graves’ disease, AITD, TSH level, TPOAb level)^30^. This was also the case when analyses were extended to single nucleotide polymorphisms (SNPs) in linkage disequilibrium (LD; r^2^>0.8). On the other hand, using the adjacent genes identified in previous GWASs and reported in GWAS catalog, we identified 14 genes that have SNPs in or near the gene body, associated with both immune cell traits (30 lead SNPs) and thyroid phenotypes (17 lead SNPs) (**Supplementary Table 4**). Two genes, *HCP5* and *MIC-A*, are associated with TPOAb-positivity and Graves’ disease as well as with three immune cell traits (two NK cell types, 16+56/314-158a+ and 16+56/CD2-314+335-337-158a+158b+, and one dendritic cell type, 11c+ (nodim)/1c-/16+/32+) (**Fig. 3b**).

We then investigated the effect of 47 SNPs associated with immune cell traits and thyroid phenotypes on gene expression in blood and thyroid tissue using eQTLs from the GTEx project and other publications^32, 33, 46^. No genetic variants previously associated with AITD or other thyroid phenotypes appeared to be also associated with the expression of CD335 receptors or its known ligands in blood and thyroid cells. On the other hand, we observed that two SNPs, rs1521 and rs3094228, associated with Graves’s diseases and TPOAb-positivity respectively fall in the gene regulatory regions of *MIC-A* and *MIC-B* genes, two ligands of CD314 (*NKG2D)*, and also increase their gene expressions in thyroid cells as well as other cell types^32, 33, 47–49^ (**Fig. 3b, Supplement Table 4, Supplement Fig. 2**). Consequently, this up-regulation of *MIC-A* and *MIC-B* gene expressions in thyrocytes could activate the cytotoxicity of NK cells and the cytokine production against thyrocytes when there is binding between NK cells and thyrocytes. Furthermore, rs1521 also reduce the expression of the *HLA-C* gene, ligand of CD158b, in the thyroid cells, as well as whole blood and other cell types^32, 33, 46^ (**Fig. 3b, Supplement Table 4, Supplement Fig. 2**). This result suggests a potential mechanism where down-regulation of HLA-C gene expression in thyrocytes caused by genetic variants could not inhibit the activation of cytotoxicity of NK cells and cytokine production when there is binding between NK cells and thyrocytes. Two of three SNPs, rs2596460 and rs252369, associated with the level of the subpopulation of NK featuring 16+56/CD2-314+335-337-158a+158b+, fell in the same locus (LD r^2^> 0.8) as the AITD-SNP, rs1521, and decrease the expression of *HLA-C* gene in the whole blood.

Overall, two SNPs, rs1521 and rs3094228, associated with respectively Graves’ disease and TPOAb-positivity, appear to alter the expression of ligands of two immunoreceptors of NK cells, CD314 and CD158b in thyrocytes, that could increase the activation of cytotoxicity of NK cells after binding with target cells (**Supplement Fig. 2**).

### AITD is associated with increased serum Caspase-2 and IL-1α abundances

In addition to the production of thyroid autoantibodies and reduced levels of IgG core fucose with a subpopulation of NK cells in AITD, we evaluated whether the abundance of 1,113 free soluble proteins in peripheral blood may be associated with AITD status (27 AITD patients versus 130 healthy controls) and TPOAb levels (155 individuals) in the TwinsUK cohort (**Supplement Table 1**) using aptamer-based multiplex protein assay (SOMAscan)^55^. As there are partial correlations between proteins (**Fig. 4a**), we identified 227 independent features using the Li and Ji’s method^45^ (Bonferroni multiple testing correction, P-value<1.9×10^-4^). Levels of three proteins were positively associated with AITD status: TSH (P-value=8.67×10^-5^; Beta=0.67; SE=0.16), Caspase-2 (CASP-2; P-value=2.72×10^-7^; Beta=1.10; SE=0.20) and Interleukin-1α (IL-1α; P-value=7.46×10^-5^; Beta=0.41; SE=0.09). We also observed a higher level of TSH on average in all patients with AITD or TPOAb-positivity compared with control individuals (euthyroidism with TPOAb-negative). We observed a significant and moderate correlation of TSH levels from two types of TSH measurement regardless of manufacturer with ones from SOMAscan (**Fig. 4b**). These indicate that the SOMAscan assay is a relevant new technology to quantify the secretion levels of soluble proteins, as previously described^56, 57^. Although Caspase-2 and IL-1α levels were associated with AITD status, Caspase-2 and IL-1α levels were not associated with TPOAb or TSH levels as continuous variables (P-value>1.9×10^-4^). However, when participants were divided into 4 categories according to TSH and TPOAb levels (**Fig. 4c**), reflecting different clinical categories (hyperthyroidism, euthyroidism/TPOAb-negative, hypothyroidism and euthyroidism/TPOAb-positive), Caspase-2 showed significantly higher mean and variance in two groups: hypothyroidism and euthyroidism/TPOAb-positive compared to euthyroidism/TPOAb-negative and hyperthyroidism (**Fig. 4d**). Participants in these two groups are likely to have underlying Hashimoto’s thyroiditis (HT). On the other hand, the variance of IL-1α was significantly larger in groups with euthyroidism/TPOAb-positive and hyperthyroidism than in the groups with euthyroidism/TPOAb-negative and hypothyroidism (**Fig. 4e**), but no significant difference for their mean. This means that on average, individuals from 4 categories have the same level of secretion of IL-1α, but there are more inter-individual variabilities in euthyroidism/TPOAb-positive and hyperthyroidism than euthyroidism/TPOAb-negative and hypothyroidism.

**Figure 4.**
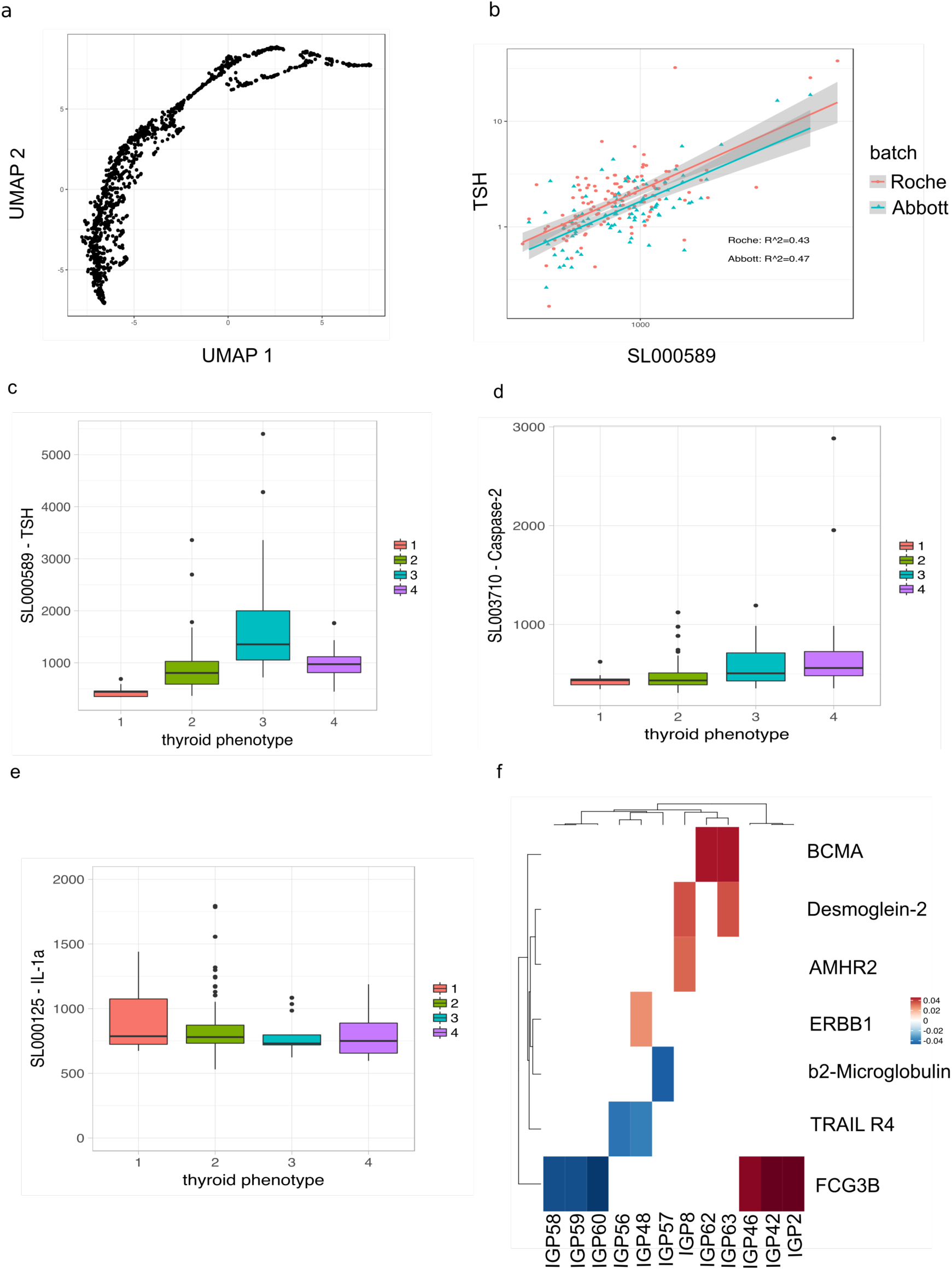
Association of circulating protein abundances with thyroid diseases and with AITD-IgG N-glycan structures. a) 1,113 circulating proteins were arranged in two dimensions based on the similarity of their secretion profiles in the serum by the dimensionality reduction technique UMAP^50^ using R package umapr^51^. b) Correlation of log10-transformed TSH measurements between two clinical FDA approved clinical immunoassays (Roche and Abbott) and SOMAscan assay in 217 individuals (122 using Roche immunoassay and 95 using Abbott immunoassay). c) Box plot of the level of circulating TSH measured by SOMAscan assay in the serum according to the group of thyroid status. d) Box plot of the level of circulating Caspase-2 measured by SOMAscan assay in the serum according to the group of TSH. e) Box plot of the level of circulating IL-1α measured by SOMAscan assay. An extreme outlier sample in the group 4 with an IL-1 α of 250,000mg/ml was discarded for the analysis. f) Heatmap of circulating protein abundances associated with AITD-IgG N-glycan structures. In fig.2c-e, participants were assigned to 4 categories according to TSH level and TPOAb status: 1=hyperthyroidism (TSH<=0.1 mIU/L; 13 individuals), 2=euthyroidism/TPOAb-negative (0.4<TSH>4 mIU/L & TPOAb < 6 IU/mL (Abbott) or TPOAb < 34 IU/mL (Roche); 196 healthy individuals), 3=hypothyroidism (TSH>=4 mIU/L; 21 individuals), and 4=euthyroidism/TPOAb-positive (0.4<TSH>4 mIU/L & TPOAb >= 6 IU/mL (Abbott) or TPOAb >= 34 IU/mL (Roche); 28 individuals).

In summary, we replicated the association of the plasma TSH levels with AITD status, and we found two novel associations of plasma protein levels of Caspase-2 and IL-1α with AITD status, but their secretion (mean and variance) seems to also depend on other factors associated with thyroid diseases such as the levels of TSH and TPOAb.

### Afucosylated IgG is associated with serum levels of several circulating proteins

We next studied the correlation between the level of secreted TSH, Caspase-2 and IL-1α proteins and IgG N-glycan trait levels in peripheral blood of 164 individuals but found no significant associations (P-value>8.3×10^-4^, Bonferroni test considering only 3 independent proteins and 20 independent IgG N-glycan traits) (**Supplement Table 6, Fig. 4f**). On the other hand, several AITD-IgG N-glycan traits appeared to be associated with other circulating proteins (P-value<3.67×10^-5^, Bonferroni test in considering only 227 independent proteins and 6 independent IgG N-glycan traits) (**Supplement Table 6, Fig. 4f**). For example, three AITD-IgG N-glycan traits (IGP2, IGP42, and IGP46) were positively associated with the circulating FCGR3B protein (FcγRIIIb or CD16b), the receptor of polymorphonuclear neutrophils (PMN), whereas three AITD-IgG N-glycan traits (IGP58, IGP59, and IGP60) were negatively associated with FCGR3B protein. Also, IGP56 and IGP48 were negatively associated with β2-microglobulin, a protein involved in the presentation of intracellular antigens through the MHC class I complex, and IGP48 was also positively associated with ERBB1 protein, also known as the epidermal growth factor receptor (EGFR), which serves as a checkpoint for cell proliferation and differentiation.

Overall, 11 AITD-IgG N-glycan traits (IGP2, IGP8, IGP42, IGP46, IGP48, IGP56, IGP58, IGP59, IGP60, IGP62, and IGP63) were associated with serum levels of 5 circulating proteins (AMHR2, BCMA, β2-microglobulin, ERBB1, and FCGR3B) in the TwinsUK cohort, but no triplet (AITD, IgG N-glycan structures, and circulating proteins) could be identified. More studies need to be performed to understand the relationships between these two components of immune responses in AITD.

### The abundance of free-soluble plasma Desmoglein-2 protein is associated with AITD genetic variants and two AITD-IgG N-glycan traits, but not significantly with AITD status

Recently, several GWASs on secreted proteins (protein quantification locus traits, pQTL) have been performed^31^, which allowed us to determine whether the secretion of proteins associated with AITD or with AITD-IgG N-glycan traits are driven by SNPs also associated with AITD. We studied whether the SNPs previously associated with plasma circulating protein abundance were also associated with at least one of thyroid phenotypes published in the GWAS catalog including GD and TPOAb-positivity^30^ and with one of 17 AITD-IgG N-glycan structures. We found no SNPs associated with one of 17 AITD-IgG N-glycan structures that are also pQTL. However, four genetic variants associated with thyroid phenotypes (rs3761959, rs7528684, rs505922, and rs3184504) are also associated in *cis* and *trans* with nine circulating protein abundances (BGAT, CHSTB, DC-SIGN, Desmoglein-2, DYR, FCRL3, GP1BA, MBL, and VCAM-1) (**Supplement Table 7**). None of these proteins were associated directly with AITD or TPOAb levels in our study. However, we found that the Desmoglein-2 protein was associated with two AITD-IgG N-glycan traits, IGP8, and IGP63^4^ (**Supplement Fig. 3)**. This protein is highly expressed in epithelial cells including thyrocytes and cardiomyocytes and plays a role in the cell-cell junction between epithelial, myocardial, and certain other cell types. It was also proposed to be a novel regulator of apoptosis^59^. Therefore, four genetic variants (rs3761959, rs7528684, rs505922, and rs3184504) are associated with nine secreted protein abundances in blood and also associated with several thyroid phenotypes. Further studies need to be performed to better understand these links (e.g., causality, pleiotropy effects).

## Discussion

The dysregulation of the immune system in AITD affects several biological structures and processes, such as antigen/antibody/Fc receptor complex formation, all driven by genetic and environmental factors. Little is known about the key players, the mechanisms of disease, and the role of genetic variants found in previous GWASs of patients with AITD. Altered biological structures and processes in AITD act in the thyroid gland, but some of them could be detected in the peripheral blood and potentially used as biomarkers. We previously identified a depletion of IgG core fucose in the peripheral blood of people with AITD, and we suggested that this feature is associated with TPOAb levels and may play a role in ADCC in patients with AITD^4^. However, ADCC, as an immunological mechanism, depends on the formation of antigen/antibody/Fc receptor complexes of substantial affinity or avidity. Thus, immune effector cells and other molecules, such as ligands and secreted proteins are also talented players in the activation of ADCC, and these were examined in this study.

We applied for the first-time an *in silico* multi-omic approach on individuals from the TwinsUK cohort to better understand the cross-talk between immune features and genetic variants in AITD. In AITD patient samples, we observed increased levels of three circulating proteins (TSH, Caspase-2, and Interleukin-1α) and a decreased level of IgG core fucosylation associated with a subpopulation of NK cells defined primarily by the expression of CD335, CD134, and CD158b receptors. Our data confirmed the previously reported association of plasma TSH level with AITD status and importantly revealed previously unknown biomarkers for AITD, which are highly associated with immunological activation functions such as ADCC, apoptosis and pro-inflammatory pathways. Several SNPs associated with AITD appear to alter the secretion of several ligands of NK immunoreceptors in thyrocytes and plasma circulating proteins, showing that genetic background may play potential roles in NK-ADCC in individuals with AITD.

The main limitation of our study is the lack of replication of our findings in other cohorts. However, to our knowledge, no other cohorts have large datasets with the same diversity of -*omics* data with AITD phenotype or TPOAb levels that would allow us to fully or even partially replicate our findings. For example, NK cells are primarily classified into only three subsets, CD56^-^, CD56^dim,^ and CD56^bright^, and not with other immunoreceptors^64^. Using high-resolution deep immunophenotyping flow cytometry, the categorization of immune cell traits performed in the TwinsUK cohort is composed of a set of several immunoreceptors. Another limitation is the absence of *in vitro* experimental evidence that could validate our findings. It is challenging to obtain thyroid tissues from healthy individuals or patients with Hashimoto’s thyroiditis, and to our knowledge, no thyroid cell lines from patients with AITD are currently available. We can, nevertheless, appreciate that different markers identified in our study contribute to concluding the presence or activation of ADCC mechanisms in AITD.

Based on the TSH and TPOAb levels, Hashimoto’s thyroiditis predominates in AITD in our cohort. This finding was also observed in several other cohorts around the world^65^. Although Hashimoto’s thyroiditis is mostly characterized by the progressive destruction of normal thyroid tissue^66^ whereas Graves’ disease is characterized by stimulatory antibodies to the thyroid-stimulating hormone receptor (TSH-R), with reduced apoptosis of thyrocytes^67^, previous studies have often not distinguished between subgroups of AITD and that could make it more difficult to interpret further findings. However, ADCC has been reported in AITD without restriction to subgroups of patients with AITD using ADCC assays with extracted antibodies from the blood of patients with the different subgroups of AITD. As expected, more patients with Hashimoto’s thyroiditis than with Graves’ disease present ADCC activities^18, 19^. Consequently, our cohort seems to represent the main mechanism in Hashimoto’s thyroiditis rather than in Graves’ disease, namely the destruction of thyroid cells.

Interestingly, two secreted proteins (Caspase-2 and IL-1α), which play a role in apoptosis and inflammatory response, were positively associated with AITD but were not linearly associated with TPOAb and TSH levels. The Caspase-2 protein mediates cellular apoptosis and plays a role in stress-induced cell death pathways and cell cycle maintenance^68^. In addition to the positive association of Caspase-2 with AITD, we observed an increase of its secretion in patients with hypothyroidism and euthyroidism/TPOAb-positive. As TPOAb has been proposed to promote ADCC against thyroid cells^4, 10, 18, 19, 69^, the Caspase-2 protein may represent a marker signifying the destruction of thyroid cells by TPOAb. The IL-1α protein, on the other hand, is a member of the interleukin 1 cytokine family and is produced mainly by activated immune cells as well epithelial and endothelial cells in response to cell injury and induces apoptosis. It is thus considered an apoptosis index of the target cell^70^. Thyroid follicular cells produce IL-1α as well as IL-6 (no significant association detected in this study), proportional to the degree of lymphoid infiltration in thyroid disorders^62^. IL-1α seems to reduce the thyroid epithelial barrier without necessary signs of general cytotoxicity^63^. In our study, the IL-1α protein was secreted more in AITD than in healthy individuals and its variance was greater in euthyroidism/TPOAb-positive and hyperthyroidism. IL-1α levels thus could be a biomarker of dysregulation in cell structures in the thyroid gland and suggest high variability in the levels of injured thyroid cells in patients with AITD. Overall, one could speculate that Caspase-2 and IL-1α could be potential biomarkers of the degree of tissue dysregulation, thyroid cell death or apoptosis and lymphoid infiltration of the thyroid gland.

Glycome-wide association studies with immune cell traits in the general population suggested that IgG core fucose levels are associated with a subpopulation of NK cells. Here, in the bloodstream of the TwinsUK cohort we observed that IgG with core fucose was positively associated with a subpopulation of NK cells with an activating NK receptor (CD314) and a differentiation receptor (CD335) profile, whereas the same IgG with core fucose was negatively associated with a subpopulation of NK cells with an inhibitory NK receptor (CD158b) and conversely between the same subpopulations of NK cells featuring CD335, CD314, CD158b immunoreceptor expression with IgG without core fucose. In the presence of IgG core fucose, three immunoreceptors, CD335, CD314, CD158b, of NK cells appeared to play a role in the activation of cytotoxicity of NK cells when they interact with their ligands on the target cells, which are thyrocytes in AITD^39–44, 71^, and this is expected to increase inflammation and susceptibility to autoimmune disease^72^.

The role of IgG core fucosylation in ADCC could thus be enhanced via a cross-talk with a subpopulation of NK cells. Previous studies showed that afucosylated antibodies have a much higher affinity (100-fold) for FcγRIIIa (CD16a) and so, enhance ADCC^73^. Moreover, a study using *in vitro* assay and human peripheral blood cells showed that ADCC via FcγRIIIa requires NK cells, but not monocytes or polymorphonuclear cells and that the activity levels of the antigen/antibody/effector cell complexes correlated only with the NK cell numbers present in the PBMCs^24^. Our associations between the level of IgG core fucose and of a subpopulation of NK cells reinforce the notion that there is a complementarity between IgG core fucose levels and NK cells, that could influence ADCC potency.

Moreover, our findings suggest that two opposing mechanisms combining NK receptors with IgG core fucose levels are working together to achieve homeostasis in NK cell activation, i.e. with IgG glycosylation as a way to fine-tune the balance between activation and inhibition, depending on the NK cell repertoire, or conversely, the NK cell repertoire may be a way to fine tune the presence of IgG. It is noteworthy that a complex balance between activating receptors, inhibitory receptors and co-receptors controls NK cell activation and cytotoxicity^74^. However, it was proposed that when antibodies specific to an antigen, such as TPOAb on TPO, cross-link NK cells with target cells, the activating stimulus from FcγRIIIa of NK cells overcomes inhibitory signals^75^. This leads to the activation of NK cells, which release cytotoxic granules containing perforin and granzymes, potentiating ADCC. Consequently, fine-tuning via the IgG glycosylation proposed here can be complementary to the presence of multiple receptors on NK cells in the activation of ADCC. This concept merits further investigation.

The role of NK cells in AITD has been previously studied, but the measurement of NK activity in the blood of individuals with AITD using *in vitro* assays produced variable and potentially contradictory results, at first view^76, 77^. For example, one study showed that an increase of NK cell activity is associated with a worsening of AITD, both in HT and GD, and suggests that NK cells might contribute to the severity of disease in autoimmune thyroid conditions^76^. On the other hand, another study observed a functional defect of a subpopulation of NK cells in the bloodstream of individuals with early-HT and proposed that this could lead to the expansion of auto-reactive T-and B-cells and so, to the pathogenesis of HT^77^. However, NK cell functions may be specific to their local tissue microenvironment and potentially also specific to the stage of disease, as NK cells may exert both cell-mediated cytotoxicity and regulatory functions^78^. Consequently, NK cells may be involved in various stages of AITD, both HT and GD, development through different roles.

Although AITD are highly heritable (55-75%)^4^, no common additive genetic variance between these different features studied here and AITD could be determined. We previously estimated the heritability of AITD, IgG N-glycan traits, and immune cell traits (**Supplement Table 8)**^4, 27^. Although GWAS signals for a large panel secreted protein levels were identified^31^, their heritability has not yet been estimated, and a previous study estimated the heritability of only 342 secreted protein abundances^58^. To tackle this gap, we estimated the proportion of genetic and environmental variance of 1,129 proteins studied in the present paper using the Structural Equation Modeling and twin structures present in the TwinsUK cohort (**Supplement Table 9)**. We found a small proportion of proteins having additive genetic variances in their heritability, which is in concordance with previous findings^58^. As the best model of heritability in AITD is only with dominant genetic variance, the shared genetic variance between AITD and proteins could not be estimated with accuracy as well as with IgG N-glycan traits and immune cell traits.

On the other hand, in our study, we identified several genetic variants previously associated with thyroid phenotypes to be also associated with the secretion of proteins and ligands of two immunoreceptors. Specifically, we found that two genetic variants, rs1521 and rs3094228, associated with GD and TPOAb-positivity, alter the expression of thyroid cell-expressed ligands, *MIC-A, MIC-B, HLA-C*, known to recognize CD134 and CD158b immunoreceptors expressed on NK cells. Thus, individuals having the variant rs1521 associated with GD have a reduction of expression of *HLA-C* gene, but an increase of expression of MIC-B in thyrocytes whereas the carriers of rs3094228 genetic variant associated with TPOAb-positivity have an increase of the expression of *MIC-B* gene in thyrocytes. Consequently, if their thyrocytes cross-talk with the subpopulation of NK cells with CD158b and CD314 immunoreceptor, they could not inhibit, but trigger the production of cytokines and cytotoxicity by these NK cells, and thus trigger ADCC. Thus, drawing from the knowledge generated in a large number of previous reports and from our present findings, our study highlights different immune features (glycan structures on antibodies, a subpopulation of NK cells, the secretion of Caspase-2 and IL-1α) as potential biomarkers of AITD status detectable in the bloodstream in addition to TSH and TPOAb levels. Moreover, if one speculates that active antibodies with low core-fucose are thyroid autoantibodies (e.g., TPOAb) and target cells are thyroid cells, we could propose that these biomarkers crosstalk in an antibody-dependent NK cell-mediated cytotoxicity-associated fashion in individuals with AITD^10, 79^. In support of this, we found that two genetic variants, rs1521 and rs3094228, associated with AITD which could participate in this crosstalk in altering the expression of ligands (*MIC-A, MIC-B*, and *HLA-C)* of NK immunoreceptors (CD314 and CD158b) in thyrocytes. We thus propose a model of positive autoreactive antibody-dependent NK cell-mediated cytotoxicity in the thyroid gland of AITD patients illustrated in **Fig. 5**^15, 18, 19, 80, 81^.

**Figure 5.**
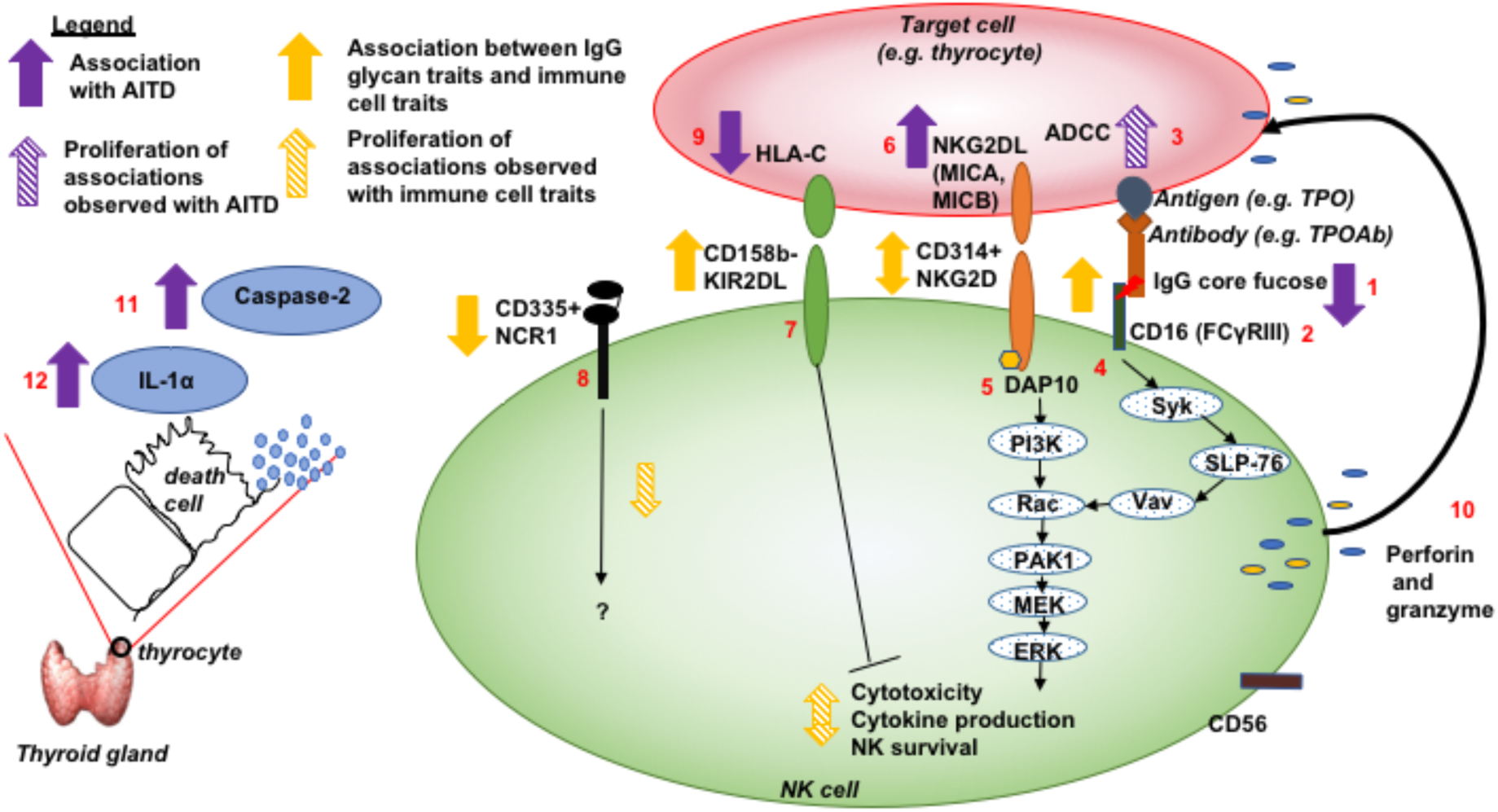
Model of different potential contributing players and their pathways activated in proposed antibody-dependent NK cell-mediated cytotoxicity in the thyroid gland of AITD patients. (1) The depletion of IgG core fucose was associated with TPOAb level and AITD status^4^. (2) The IgG N-glycan traits associated with AITD were also associated with a subpopulation of NK cells in our current study; for example, the depletion of IgG core fucose is associated positively with NK cells with the patterns of co-receptors CD335- or CD335-CD158b+CD314+. (3) Previous studies showed that a-fucosylated antibodies had increased affinity for binding to CD16 (FcγRIIIa), cell receptors of NK cells, and to enhance ADCC^22–24, 26^ via (4) protein tyrosine kinase-dependent pathways, through crosstalk with (5) NKG2D receptor (CD314)^60, 61^. (6) Two SNPs, rs3094228 and rs1521, were associated with GD and TPOAb-positivity^47–49^ and fall in gene regulatory regions of the *MIC-A* and *MIC-B* genes and increase their expression in thyroid cells^32^. These two genes encode heavily glycosylated proteins that are ligands for the NKG2D type II receptor (CD314). (7) The KIR2DL (CD158b) receptor is known to regulate the cytotoxicity of NK cells by unknown pathways, whereas (8) the NCR1 (CD335) receptor can contribute to the increased potency of activated NK cells to mediate cell lysis by unknown pathway^39, 40^. (9) The SNP, rs1521 associated with GD^47^, is also shown to reduce the expression of *HLA-C* gene, producing the ligand of CD158b, in thyroid cells^32, 33, 43, 44^. (10) All together (the binding of NK cells with target cells through antibodies and their ligands), these lead to the activation of NK cells, which release cytotoxic granules containing perforin and granzymes. This release mediates ADCC of target cells (3), which are thyrocytes in AITD. Also, (11) a positive association between the circulation abundance of Caspase-2 protein and AITD were found in this study that could be associated with the destruction of thyrocytes. (12) A positive correlation of circulating abundance of IL-1α with AITD was also found in the bloodstream that could be a marker of lymphocyte infiltration in the thyroid gland of individuals with AITD, and thus of inflammation^62, 63^.

We also speculate that targeting TPOAb, perhaps only by their glycosylation^82^, or a subpopulation of NK cells in patients with HT may lead to a deceleration in the destruction of the thyroid gland and could lead to potential complementary targets for drugs in the treatment of hypothyroidism and HT. On the other hand, the combination of afucosylated TPOAb with the activation of a specific subpopulation of NK cells could trigger the lysis of thyroid cells and could also be a new potential treatment in disseminated radioactive iodine-resistant differentiated thyroid cancer. A previous *in vitro* study^69^ tested the hypothesis that TPOAb from patients with GD could be used as thyroid cancer immunotherapy and showed potential to destroy thyroid cells, more precisely thyroid cancer cells that express cell surface TPO proteins, by ADCC or CDC involving monocytes or NK cells as effector cells. They observed that the deglycosylation of TPOAb reduced the binding to FcγRs and thus inhibited the apoptosis of thyroid cancer cells via CDC and cytotoxicity via ADCC, showing the importance of glycosylation on TPOAb in the destruction of thyroid cancer cells. We suggest that the production of afucosylated thyroid autoantibodies with the activation of specific subpopulations of NK cells could enhance the cytotoxicity activities of TPOAb on cancer thyrocytes. Advanced new technologies in glycosylation engineering^82–84^ and in immunotherapy such as the activation of specific subpopulations of NK cells^85–89^ in tumors could help to validate this hypothesis.

## Materials and Methods

### Study Sample

The samples for immune cell traits, glycosylation, proteomics, and GWAS for immune cell traits consisted of twins from the UK Adult Twin Registry (TwinsUK cohort). The TwinsUK cohort is comprised of approximately 12,000 monozygotic and dizygotic twins unselected for any particular disease or trait. The cohort is from Northern European/UK ancestry and has been shown to be representative of singleton populations and the UK population in general^90, 91^. The project was approved by the local Ethics Committee, and informed consent was obtained from all participants. The TwinsUK cohort for each omics is described in **the Supplementary Table 1**. TwinsUK glycomic, immunological, transcriptomic, proteomic, genetic data and GWAS results are publicly available upon request on the department website (http://www.twinsuk.ac.uk/data-access/accessmanagement/).

### Detection of TSH and TPOAb and Definition of AITD

The study was performed using a clinical AITD definition and TPOAb as a threshold trait as we do not have clinical diagnostic data related to AITD confirmed by a clinician. Individuals were considered to have AITD if they had a positive TPOAb titer (3 times more than the threshold set by the manufacturer [18 IU/mL for Abbott assay and 100 IU/mL for Roche assay]) or had a TSH level more than 10 mIU/L. On the other hand, we considered individuals as controls if they had a TSH level in the normal range and a negative TPOAb titer, with no previous clinical diagnosis of thyroid disease and not treated with thyroid medications or steroids. Individuals with a history of thyroid cancer or thyroid surgery were excluded.

Sera to assess TPOAb and TSH levels were collected between February 1994 and May 2007 and kept frozen at −80°C until use. Quantitative determination of TSH and TPOAb (only IgG class) levels was performed on the sera either by a chemiluminescent microparticle immunoassay (CMIA) [ARCHITECT® Anti-TPO or TSH (ABBOTT Diagnostics Division, Wiesbaden, Germany, 2005)] (TPOAb titer>6 mIU/L considered positive; reference range for TSH level 0.4-4.0 mIU/L) or by an electrochemiluminescence immunoassay “ECLIA” [Elecsys and Cobas e analyzers, (Roche Diagnostics, Indianapolis, lN, USA, 2010)] (TPOAb titer>34 IU/mL considered positive; reference range for TSH 0.4-4.0 mIU/L).

### Detection of IgG glycosylation profiling for discovery

Plasma for the analysis of IgG glycosylation was collected between 1997 and 2013 in 2,279 individuals from the TwinsUK cohort. The IgG glycosylation profiling was performed on total IgGs glycome from plasma (combined Fc and Fab glycans and all IgG subclasses) in Genos Glycoscience Research Laboratory, Croatia using UPLC analysis of 2AB-labelled glycans. The protocol, as well as the data pre-processing and normalization of data in the TwinsUK cohort, was previously described^4^. Briefly, using UPLC analysis of 2AB-labelled glycans, the chromatograms were all separated in the same manner into 24 peaks, and the amount of glycans in each peak was expressed as a percentage of the total integrated area. One glycan was excluded before any transformation and standardization of data because of its co-elution with a contaminant that significantly affected its values in some samples whereas two glycan peaks (GP) GP20 and GP21 (Zagreb code) were combined into a single trait called GP2021 (Zagreb code) because of difficulty in distinguishing between these peaks in some samples. A global normalization and natural logarithm transformation were applied to 22 directly measured glycan structures. As many of these structures share the same structural features (galactose, sialic acid, core-fucose, bisecting *N*-acetylglucosamine (GlcNAc)), 55 additional derived traits were calculated that average these features across multiple glycans from the 22 normalized and non-transformed directly measured glycans. Technical confounders (batch and run-day effects) were addressed using R package ComBat. The 22 directly measured glycans and 55 derived glycan traits were centered and scaled to have a mean of 0 and standard deviation (SD) of 1. Samples being more than 6 SD from the mean were considered as outliers and excluded from the analysis.

### Detection of immune cell traits

Plasma samples for assessment of 78,000 immune traits have collected between 2010 and 2012 in 669 female participants from the UK Adults Twin Register, TwinsUK (497 for a discovery cohort and 172 for a replication cohort) using high-resolution deep immunophenotyping flow cytometry analysis with a protocol described previously^28^. 78,000 cell-types captured by 7 distinct 14-color immunophenotyping panels were detected and described immune cell subset frequencies (CSF) and immune cell-surface protein expression levels (SPELs). After quality control removing immune cell traits that appeared as poor reproducibility or out of range, 23,485 immune cell traits from 497 individuals of the discovery cohort were analyzed in this study. For the current analysis, only 374 twins have immune cell traits data and TPOAb level detected by Roche immunoassay and 245 individuals in a case-control study in combining Roche and Abbott assays (204 controls and 41 AITD). Immune traits were quantile normalized residuals of a linear mixed effect model where age was included as fixed effects, and the batches were considered as random effects.

### Detection of protein profiling in plasma

1,129 proteins were measured in 2013 on plasma samples collected between 2004 and 2011 from 211 female twins from the TwinsUK Adult twin registry using SOMAscan v2 (SomaLogic Inc, Boulder, CO). The protocol was described in a previous paper^92, 93^. Briefly, hemolyzed samples were first excluded. Proteins were then measured using a SOMAmer-based capture array called “SOMAscan.” Quality control was performed at the sample and SOMAmer level and involves the use of control SOMAmers on the microarray and calibration samples. At the sample level, hybridization controls on the microarray are used to monitor sample-by-sample variability in hybridization, while the median signal over all SOMAmers is used to monitor overall technical variability. The resulting hybridization scale factor and median scale factor are used to normalize data across samples. The acceptance criteria for these values are 0.4–2.5, based on historical trends in these values. Somamer-by-somamer calibration occurs through the repeated measurement of calibration samples; these samples are of the same matrix as the study samples and are used to monitor repeatability and batch to batch variability. Historical values for these calibrator samples for each SOMAmer are used to generate a calibration scale factor. The acceptance criteria for calibrator scale factors is that 95% of SOMAmers must have a calibration scale factor within ±0.4 of the median. For the current analysis, only 1,113 proteins were then studied.

### Statistical Methods

All statistical analyses were run using R version 3.2.3. Linear mixed effect models were done in using R lme of package lme4^97^, and linear models were done in using R function lm of package stat.

#### Determination of effective number of independent tests for different omic data

Due to high and partial correlations within both glycans and immune cell traits, we decided to use the equation 5 proposed by Li & Ji (2005)^45^ to define an effective number (M_eff_) of independent tests. We then used this number to define the effective Bonferroni P-value threshold such as 0.05/M_eff_ instead of 0.05/M, with M the actual number of tests. 20 independent tests were estimated for 76 glycans. Consequently, to account for multiple testing in the discovery cohort, we present results surpassing a conservative Bonferroni correction assuming 20 independent tests, thus giving a significant threshold of (P-value <2.5×10^-3^=0.05/20). 1,357 independent tests were estimated for 23,485 immune cell traits, thus giving a significant threshold of 3.68×10^-5^ (0.05/1,357). 227 independent tests were estimated among 1,113 proteins (P<0.05/227=1.9×10-4).

#### Association studies between omics features and thyroid phenotypes

To examine whether one of the 17 AITD-IgG N-glycan traits was significantly associated with one of the 23,485 immune cell traits, we compared the fitted model in equation (2) with a model that did not include the residual of glycan in equation (1):

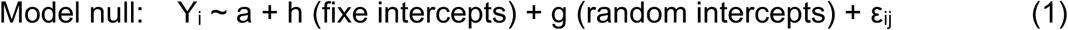

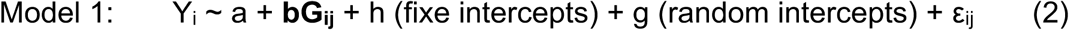

Where Y*_i_* represents the quantification of immune cell traits for individual *i* and Gij is glycan structure of type *j* among 75 *N-*glycans for the same individual i. If biological covariates (age, sex) have not been adjusted before association analysis, they have been added in the model. A random intercept was added only in the discovery cohort in order to model the family-relatedness.

To examine whether an immune cell trait was significantly associated with TPOAb level and AITD status, we compared the fitted model in equation (2) with a model that did not include the immune cell traits in equation (1) where Gij become the immune cell trait of type j among 23,485 in discovery cohort for the same individual i. For the discovery and replication cohorts in TwinsUK, we added a random intercept in order to model the family-relatedness.

To examine whether one of the 1,113 protein was significantly associated with TPOAb level and AITD status, we compared the fitted model in equation (2) with a model that did not include the protein in equation (1): where Gij become the protein of type j among 1,129 in discovery cohort for the same individual i. We added a random intercept in order to model the family-relatedness.

To examine whether one of 1,113 proteins was significantly associated with one of 17 significant glycans, we compared the fitted model in equation (2) with a model that did not include the protein in equation (1): where Gij become the protein of type j among 1,129 in discovery cohort for the same individual i. We added a random intercept in order to model the family-relatedness.

#### Genome-wide Association Analysis on IgG N-glycan traits, immune cell traits, proteins, gene expressions in several tissues, and thyroid phenotypes

To define the list of SNPs associated with glycosylation profiles regardless of specific phenotypes in the TwinsUK cohort, we ran analyses using GenABEL software package^98^, which is designed for GWAS analysis of family-based data by incorporating pairwise kinship matrix calculated using genotyping data in the polygenic model to correct relatedness and hidden population stratification. We selected SNPs for each IgG N-glycan traits that have a P-value under GWAS P-value threshold (P-value < 5×10^-8^). Moreover, we added the list of SNPs previously defined^29, 99^.

To define the list of SNPs associated with immune cell traits regardless to any specific phenotypes in the TwinsUK cohort, we ran extracted SNPs for each immune cell traits that have a P-value under GWAS P-value threshold (P-value < 5×10^-8^) from previous published GWASs on these immune cell traits^27, 28^.

To define the list of SNPs associated with protein abundance found in this study, we extracted the significant SNPs reported in INTERVAL project^31^.

To define the list of SNPs associated with gene expression (eQTL), we extracted the eQTLs reported significant by GTEx and previous papers present in HaploReg V4.1^31,32^.

To define the list of SNPs associated with AITD and thyroid functions, we selected to SNPs listed in the NHGRI GWAS catalog^30^ with words “thyroid” or “Graves” or “Hashimoto.”

#### Determination of shared SNPs and genes between IgG N-glycan traits, immune cell traits, proteins abundance, and thyroid functions and diseases

To examine whether glycans, immune cell traits, and thyroid functions and diseases shared SNPs or genes, we compared the list of SNPs from GWASs done on TwinsUK data and from the NHGRI GWAS catalog (cf. above). As SNPs detected by GWASs could be lead SNPs and not necessarily causal SNPs^100^, we extended the list of SNPs to other variants in linkage disequilibrium (LD) with an r² threshold of 0.8 from 1000G Phase 1 European population. Using HaploReg V4.1^101^, we extracted the closest annotated genes and transcript from GENCODE.

#### Heritability analysis for proteins

Using twin data and ADCE models (additive genetics (A), dominante genetics (D), shared environment (C) and non-shared environment (E)), heritability of glycosylation structures, immune cell traits and AITD were estimated using the R package called mets that allows us to run the analysis with monozygotic and dizygotic twins as well as unrelated individuals. The significance of variance components A, D, and C was assessed by dropping each component sequentially from the full model (ADCE) and comparing the sub-model fit to the full model. Sub-models were compared to full models by hierarchical χ^2^ tests. The difference between log-likelihood values between sub-model and full model is asymptotically distributed as χ^2^ with degrees of freedom (df) equal to the difference in df of sub-model and the full model. A statistical indicator of goodness-of-fit is the Akaike information criterion (AIC), computed as χ^2^ −2df; sub-models are accepted as the best-fitting model if there is no significant loss of fit when a latent variable (A, C, D, or E) is fixed to equal zero. When two sub-models have the same AIC compared to the full model, we decide to keep the model the most likely (with additive genetic variance) or with the lowest P-value for different components.

#### Visualization

Heatmaps were created in using R package ComplexHeatmap, and we visualized only beta values having significant associations from linear mixed models (Fig. 2a and 4f). Correlation between 51 immune cell traits and between 17 AITD-IgG N-glycans (Fig 2b and c) were created with R package corrplot. Boxplots and scatter plot were created in using R package ggplot2 (Fig 4b-e).

## Acknowledgment

- TwinsUK. The study was funded by the Wellcome Trust; European Community’s Seventh Framework Programme (FP7/2007-2013). The study also receives support from the National Institute for Health Research (NIHR)-funded BioResource, Clinical Research Facility and Biomedical Research Centre based at Guy’s and St Thomas’ NHS Foundation Trust in partnership with King’s College London and the Australian National Health and Medical Research Council (PG 1087407).
- IgG N-glycan analysis was performed in Genos and partly supported by the European Community’s Seventh Framework Programme grants HighGlycan (contract #278535), MIMOmics (contract #305280), HTP-GlycoMet (contract #324400) and IntegraLife (contract #315997).
- SNP Genotyping was performed by The Wellcome Trust Sanger Institute and National Eye Institute via NIH/CIDR.
- This study represents independent research partly funded by the National Institute for Health Research (NIHR) Biomedical Research Centre at South London and Maudsley NHS Foundation Trust and King’s College London. The views expressed are those of the authors and not necessarily those of the NHS, the NIHR or the Department of Health. This study was supported by research (RJBD) at the National Institute for Health Research University College London Hospitals Biomedical Research Centre, and by awards establishing the Farr Institute of Health Informatics Research at UCLPartners, from the Medical Research Council, Arthritis Research UK, British Heart Foundation, Cancer Research UK, Chief Scientist Office, Economic and Social Research Council, Engineering and Physical Sciences Research Council, National Institute for Health Research, National Institute for Social Care and Health Research, and Wellcome Trust (grant MR/K006584/1). SJK is supported by an MRC Career Development Award in Biostatistics (MR/L011859/1).
- The authors acknowledge support by Breast Cancer Now (147); and the Medical Research Council (MR/L023091/1).

---

**Supplement Figure 1:**
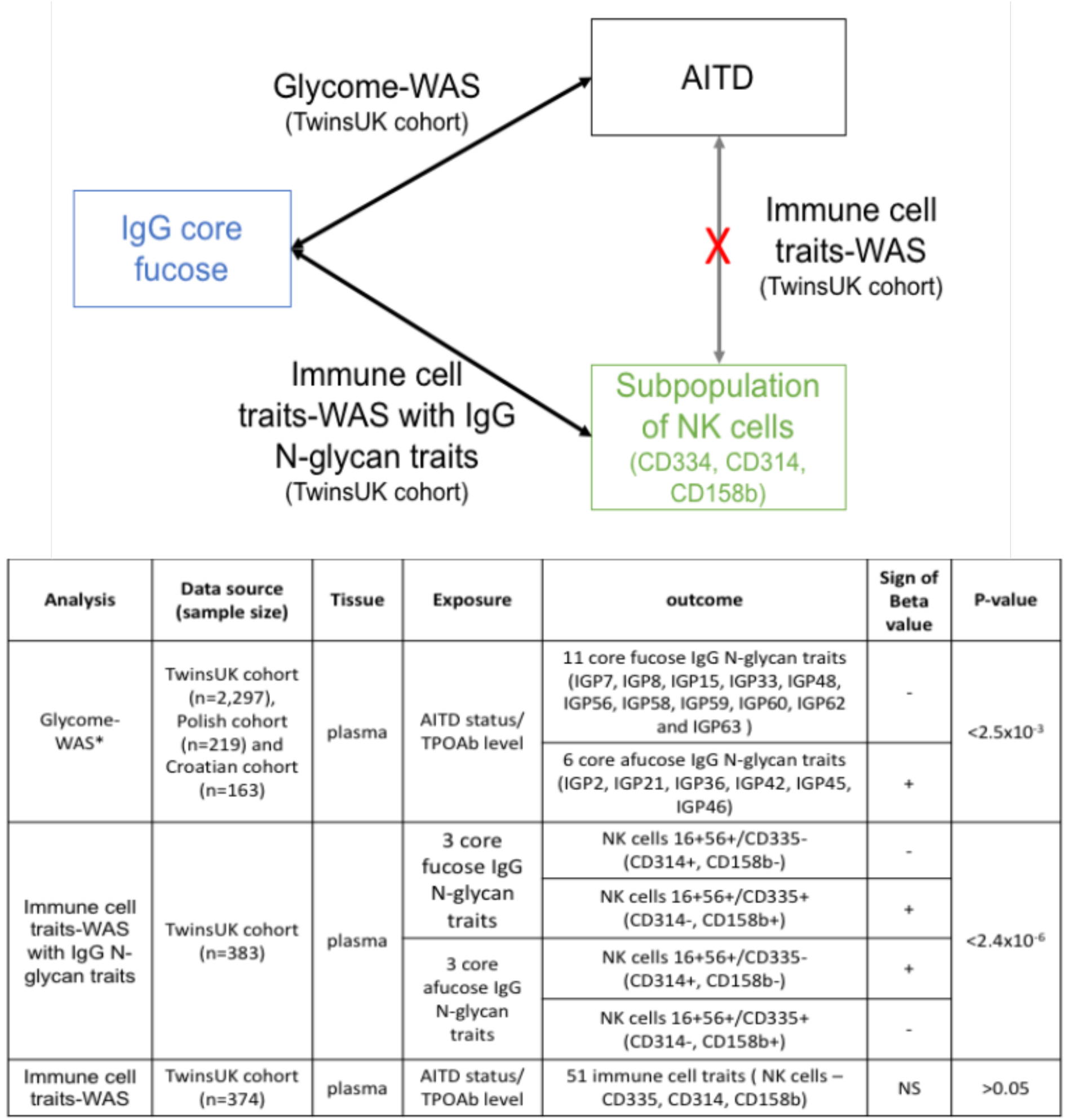
Overview of associations observed between IgG core-fucose, a subpopulation of NK cells and AITD status in the TwinsUK cohort. *Glycome-wide association studies of AITD and TPOAb levels were previously performed^4^.

**Supplement Figure 2:**
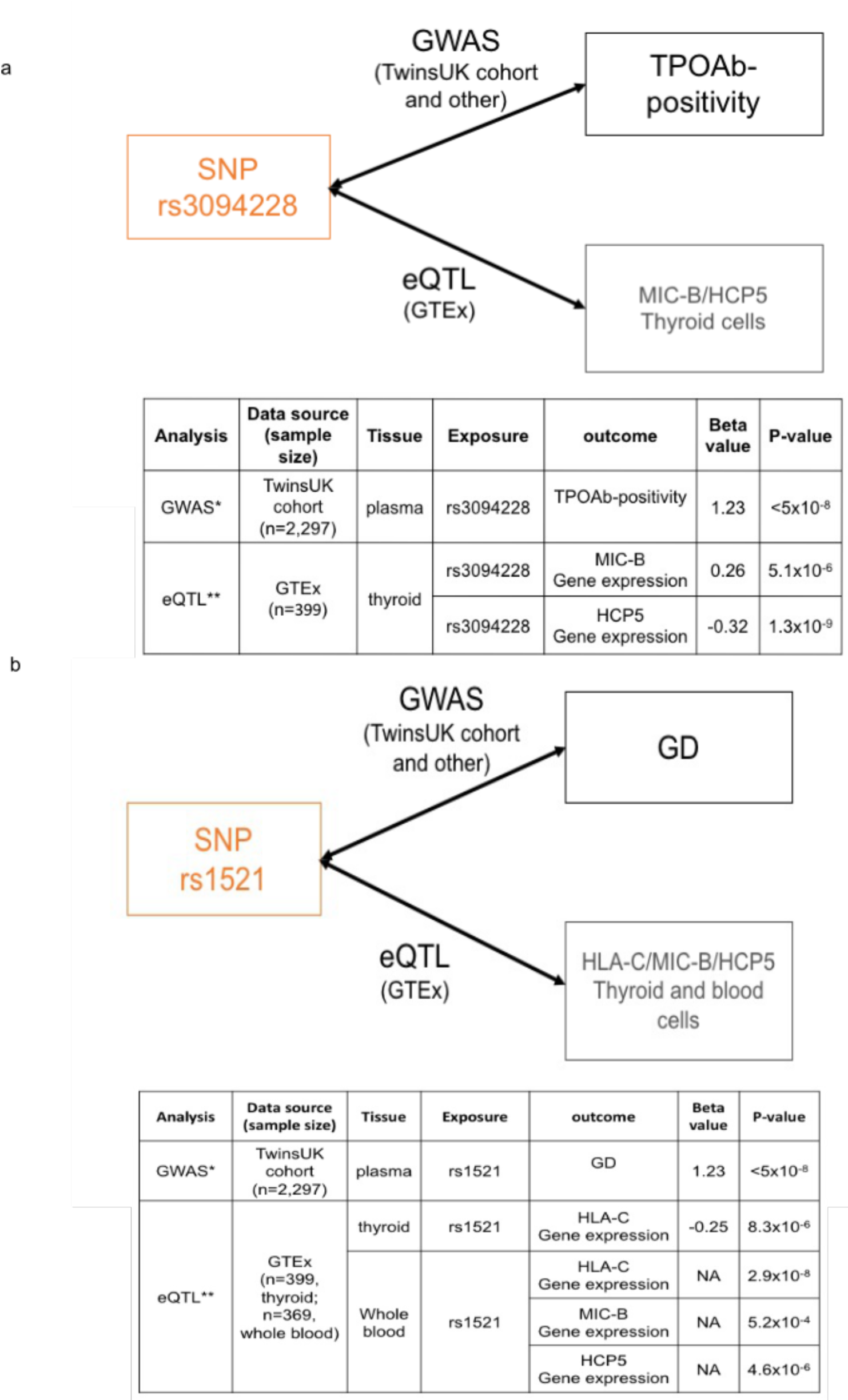
Overview of associations between AITD-SNP and eQTL in thyroid and blood cells. *Genome-wide association studies of AITD and TPOAb-positivity were previously performed, and the findings are available via GWAS catalog^27, 30^ whereas **eQTLs come from GTEx project^31, 32^.

**Supplement Figure 3:**
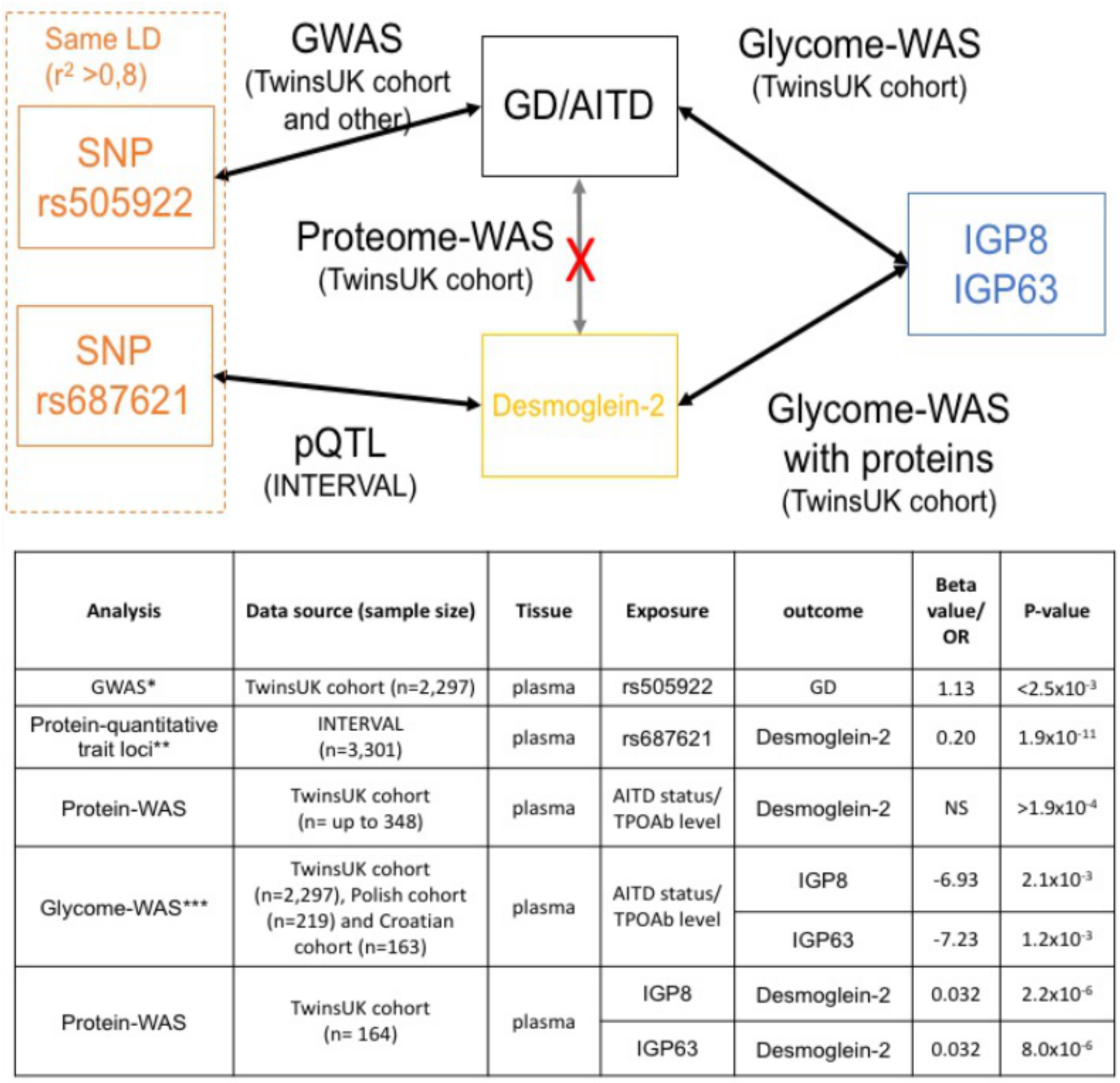
Overview of multi-omic findings associated with Desmoglein-2 in individuals with AITD status and general population. We highlighted a locus with high LD having SNPs and two IgG glycan traits that are both associated with GD and the abundance of secreted plasma Desmoglein-2 in plasma. However, no direct association of AITD status with the abundance of secreted plasma Desmoglein-2. We previously performed glycome-wide association studies of AITD and TPOAb levels^4^. Genome-wide association studies of AITD and TPOAb-positivity were previously performed, and the findings are available via GWAS catalog^27, 30^ whereas pQTLs come from INTERVAL project^31^. IGP8 = the percentage of FA2[3]G1 glycan in total IgG glycans. IGP63 = The percentage of fucosylation (without bisecting GlcNAc) of agalactosylated structures.

## Notes

**Conflict of Interests Statement:** GL is the founder and CEO of Genos Ltd, a private research organization that specializes in high-throughput glycomic analysis and has several patents in this field. MP are employees of Genos Ltd.

